# Low-cost, open-access refractive index mapping of algal cells using the transport of intensity equation

**DOI:** 10.1101/640755

**Authors:** Stephen Grant, Kyle Richford, Heidi Burdett, David McKee, Brian R. Patton

**Affiliations:** Department of Physics and SUPA, University of Strathclyde, Glasgow G4 0NG, United Kingdom; Lyell Centre for Earth and Marine Science and Technology, Edinburgh, UK, EH14 4AS; School of Energy, Geoscience, Infrastructure and Society, Heriot-Watt University, Edinburgh, EH14 4AP

## Abstract

Phase contrast microscopy allows stain free imaging of transparent biological samples. One technique, using the transport of intensity equation (TIE), can be performed without dedicated hardware by simply processing pairs of images taken at known spacings within the sample. The resulting TIE images are quantitative phase maps of unstained biological samples. Therefore, spatially resolved refractive index information can also be determined.

Using low-cost, open-source hardware, we applied the TIE to living algal cells to measure their refractive index. We obtained refractive index values that were repeatable within species and differed by distinct amounts depending on the species being measured. We suggest TIE imaging as a method of discrimination between different algal species and, potentially, non-biological materials, based on refractive index. Potential applications in biogeochemical modelling and climate sciences are suggested.

## 1 Introduction

Phytoplankton – free-living unicellular algae found throughout the world’s marine and fresh-water bodies – play a pivotal role in aquatic ecosystem structure and biogeochemistry. Despite representing just 1% of the Earth’s photosynthetic biomass, phytoplankton contribute an estimated 45% of global primary production (the production of organic compounds from carbon dioxide)[1]. On evolutionary timescales, this primary production led to an increase in atmospheric oxygen concentrations, resulting in the oxygen-rich, carbon dioxide-poor atmospheric composition. This was facilitated by their role in the global carbon cycle via biomass production [2], which has also led to a proportional balance of nutrients (including nitrogen, phosphorus and trace elements) between seawater and biomass – the so-called ‘Redfield ratio’ [3]. Calcifying phytoplankton, such as the coccolithophores, play additional roles in the marine carbonate cycle via the production of ‘coccoliths’ - calcium carbonate platelets which surround each cell [4]. Under bloom conditions, coccolithophores may contribute up to 40% of oceanic primary production and phytoplankton biomass [5]. Sinking of coccoliths that have been shed or when an algal cell dies results in a ‘rain’ of calcium carbonate particles through the water column, creating a transport pathway for organic and inorganic carbon from the sea surface to the seafloor [6]. The accumulation of biologically-produced calcium carbonate on the seafloor (from coccolithophores as well as other calcifying marine organisms) represents the world’s largest geological sink of carbon [4]. Despite these wide-ranging roles in aquatic primary production and biogeochemistry, site-specific community composition and species-specific physiological variability means that precise quantification of these roles remains poorly constrained in both marine and freshwater environments. Recent advances in measurement devices (e.g. flow cytometers) and autonomous sampling platforms (e.g. ocean gliders) have somewhat enabled this to be overcome, but remain hindered by high sensor costs, preventing replicated or networked datasets.

A key aim for monitoring marine biogeochemistry on global scales is to be able to relate ocean colour remote sensing signals to algal taxonomic and physiological properties [7]. Whilst a significant volume of work exists attempting to derive algal functional type from remote sensing data, this effort is impeded by difficulties in the forward modelling of bulk inherent optical properties (IOPs e.g. absorption and scattering) for algal cells. There is currently significant debate about the ability of simple Mie theory modelling, which assumes uniform spherical particles, to adequately capture the optical characteristics of complex naturally occurring particles such as phytoplankton and mineral particles. Recent work [8–10] has shown how flow cytometry can be used to establish size and bulk refractive index distributions for natural particle populations, and that these can be used to predict bulk optical and biogeochemical properties. This study used simple Mie theory, but required extrapolation of data beyond the range of observations in order to achieve closure with bulk IOPs. It has also been suggested [11] that internal structures within these particle types may be a significant contributor to the bulk optical properties. This study achieved similar levels of closure with bulk IOPs without extrapolation PSDs, but instead used a layered sphere model for optical scattering and made assumptions about layer thickness and refractive index distributions within the particles. Both approaches are reliant upon availability of relatively expensive equipment and are limited by relatively poor access to refractive index data for the particle population. There is a significant need to develop new tools to measure both the physical dimensions and bulk refractive index of these particle types, with mapping refractive index distributions across particles a highly desirable end-goal.

We propose a low cost, open access, easy to use device using microscopy techniques which would allow for morphological distinction of algal species in an adaptable, expandable platform designed to allow multiple measurement techniques to be added in a modular fashion. This small device could be incorporated into flow cytometry systems or deployed in addition to existing tools on autonomous underwater vehicles, drones, or as part of a water treatment facility monitoring system among other uses. This paper details one such proposed technique: the implementation of quantitative phase microscopy which enables refractive index extraction.

Phase contrast microscopy allows stain free imaging of transparent biological samples and reveals additional information and detail regarding fine structure when compared to bright field microscopy. Typical phase microscopy methods require the use of expensive optical elements or complex structured light schemes [12–17] which can be prohibitively expensive for many in educational settings or developing nations.

The transport of intensity equation (TIE), first described by Teague in 1983 [18], allows for phase microscopy to be carried out using a much simpler, and more affordable, instrumentation as it is a wholly computational, post processing technique.

The aim of this work was to assess the viability of using the TIE to determine the refractive index values of different strains of algae to allow for discrimination between them.

## 2 Transport of Intensity Equation

The TIE takes advantage of the relationship between the intensity evolution along the optical axis of a propagating wave and the wave’s phase in a volume confined within discrete longitudinal steps. When an optical inhomogeneity (such as an algal cell), which affects the phase of a propagating wave, is introduced, the TIE can be used to determine the effect the resulting phase disturbance would have on the intensity of the wave after propagating through the disturbance.

Conversely if the intensity is known along the propagation direction of the wave, then any phase objects which modulate the wave’s intensity can be identified using the TIE and the phase information at the desired focal plane can be reconstructed. For sufficiently transparent samples only two images along the propagation direction of the wave are needed to obtain the intensity gradient and reconstruct the phase in the focal plane as noted in [19]. However, in this case, in line with [20] and for completeness, three images are used; one depicting the object at the focal plane of interest and two defocused images at symmetrical distances from the focal plane along the propagation path.

The calculation is carried out using the three aforementioned images, *I*_1_, *I*_2_, and *I*_3_, where *I*_2_ is the image in the focal plane and *I*_1_ and *I*_3_ are taken a distance Δ*z* away. The TIE equation is as follows:

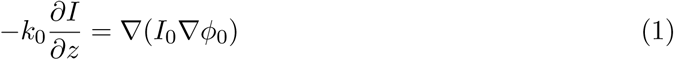

where *I*_0_ is the intensity at the focal plane, *ϕ* is the phase, and 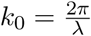 is the wavenumber. Note that *k*_0_ in this instance is a general expression with the refractive index of the medium being introduced in Eq. 4 as a scaling factor. The intensity differential, *∂*I/*∂*z is measured directly and calculated empirically from the recorded images. The introduction of an auxiliary function *ψ* allows for the conversion of Eq. 1 into a Poisson’s equation. The auxiliary function is defined as:

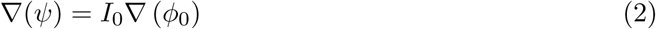

Simplifying Eq. 1, and substituting in Eq. 2, gives:

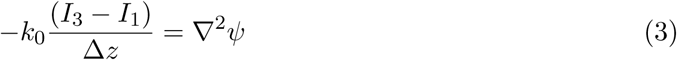

Δ*z* is the experimental distance between the two out of focus images and the focal plane. (*I*_3_-*I*_1_) is the differential between the two out of focus intensity images.

As noted in [21], in the case of uniform illumination across the sample, the phase map can be obtained by taking the Fourier transform of Eq. 3, re-weighting its Fourier coefficients in the frequency domain, and finally performing an inverse Fourier transform to obtain the phase map in the spatial domain.

Practically this means once the Fourier transform is calculated based on the empirically measured left hand side of Eq. 3, the Fourier coefficients are re-weighted in accordance with equation (3) in [20], which in turn comes from equation (11) in [21]. The inverse Fourier transform is then carried out returning the now processed auxiliary function, *ψ*, which can be integrated to give the phase, *ϕ*.

The limiting factor on the accuracy of the TIE, and subsequent RI, measurements was the accuracy of the spatial offset, Δ*z*.

Complete calculation steps, along with logic to mathematically justify them, are available in the annotated python code available in the associated project repository.

## 3 Refractive index extraction

As noted by [19], the ‘natural’ quantity returned by the TIE is Optical Path Length (OPL). This can be obtained from the *ϕ* value obtained above by using Eq. 4. This relationship is useful in that it provides a method of determining the optical path length from the phase map obtained from the TIE.

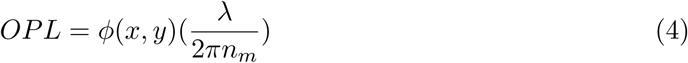

with the refractive index of the sample, *n*_*m*_. However we know experimentally that the physical path length being imaged is defined by the spacing of the two out of focus images, namely (2Δz). Therefore values which deviate from this boundary condition correspond to changes in optical path length due to changes in refractive index rather than changes in physical path length. By using:

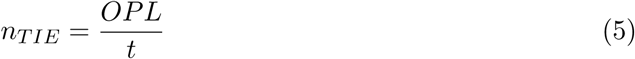

we can obtain the refractive index of the sample, *n*_*s*_, in terms of the known physical path length, *t* (equivalent to 2Δz), and the optical path length obtained from the TIE. *n*_*TIE*_ is a relative value equal to *n*_*s*_/*n*_*m*_ and returns an effective sample refractive index value, *n*_*s*_, relative to the background medium refractive index, *n*_*m*_, in this case water (*n*_*m*_ = 1.34).

## 4 Data acquisition

We used two classes of system for the experiments performed in this paper, as shown in figure 1. To test the reliability and demonstrate the effectiveness of the TIE method, the first class of instrument consisted of low cost components mounted on an optical table. We used the microscope in this configuration to perform a thorough set of measurements that are described fully in the results section. We refer to this system as the lab-prototype due to the requirement for mounting on the optical table which provides mechanical stability and minimises vibrations which may have adverse effects on optical alignment and imaging.

**Figure 1:**
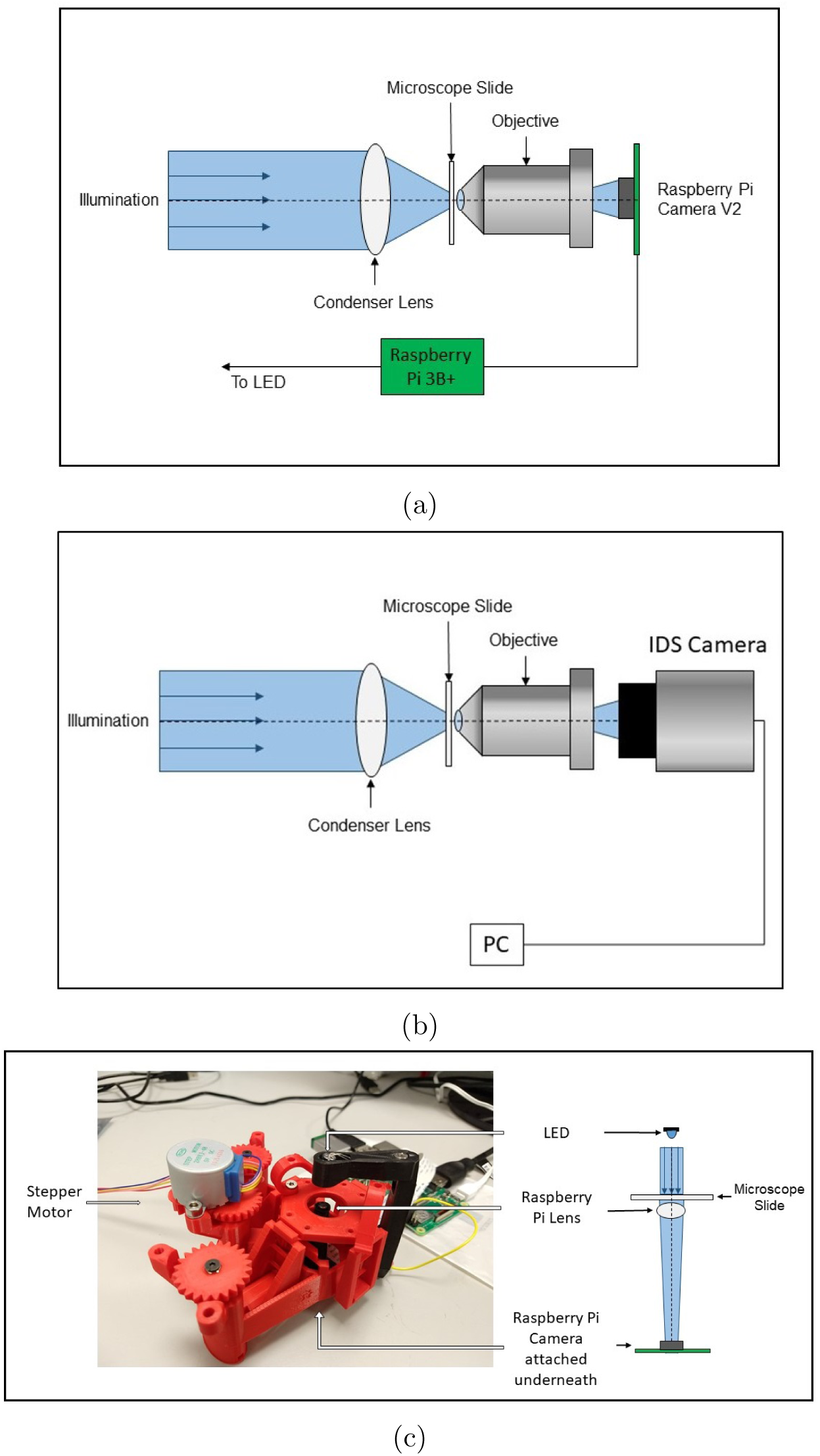
Experimental Setup showing (a) Raspberry Pi, and (b) IDS configurations lab prototypes. (c) shows the openflexure microscope with attached stepper motor. An LED is housed in the illumination arm (black piece), the Raspberry Pi Camera lens can be seen just below the center of the stage, and the camera itself is mounted on the base of the unit. Ray diagrams in all schematics are illustrative only.

In order to show how the components used in the lab-prototype could be used in a device more suitable for field deployment and that has even lower infrastructure requirements, we also performed a subset of the measurements using an open source 3D printable microscope designed by Water Scope [22]. This microscope uses a Raspberry Pi camera V2 with the camera’s lens removed, reversed and mounted at a greater distance from the sensor than in conventional use. The result is a medium numerical aperture system (≈ 0.25) that can be used in place of the camera and microscope objective systems used in the lab-prototype. We refer to this instrument as the ‘water scope’ in this paper. The water scope microscope used in this work ((c) in figure 1) was printed using an Ultimaker S5 printer and polylactic acid material. A small sample of measurements on the same algal strains were captured using a green LED for illumination. Rather than repeating all the measurements performed with the lab-prototype, we demonstrate that it is equally effective once some initial characterisation is performed. The results section for the water scope discusses the calibration and characterisation in more detail.

Figure 1 (a) and (b) show the lab-prototype apparatus. A single high brightness blue light emitting diode was used as a light source. A low cost Thorlabs condenser (ACL25416U: f=16 mm, NA=0.79) was used as the microscope condenser lens. The aqueous samples were mounted on microscope slides with an ≈250 *µ*m thick piece of electrical tape acting as a gasket to hold the liquid in place. The sample was then held vertically in a 3D printed microscope slide holder (designed by cfmccormick and licensed under CC BY-SA 3.0). This holder was mounted on a Thorlabs translation stage to allow for control of the sample position relative to the objective focal plane. A Newport objective (M60-X: m=60x, NA=0.85) was used to image the sample directly. The numerical aperture mismatch between the condenser and objective led to a reduction in possible best resolution of the objective lens in the system, i.e effectively reducing the numerical aperture of the objective to 0.79 by having less available light from the condenser.

The numerical aperture of the illumination system was estimated at 0.61 based on the condenser lens and LED used. The phase in the focal plane of the system, *ϕ*_0_, where we define the focal plane as being within the Rayleigh range of the excitation beam, should be consistent. The Rayleigh range was found to be ≈0.6 mm due to the large incident beam spot size, much larger than the step sizes of ≈10 *µ*m used when selecting planes for the TIE calculation. The system comprising the condenser lens and objective resulted in a resolution of ≈300 nm, and a depth of field of ≈1.3 *µ*m.

Two versions of the lab-prototype were tested, the sole difference between them being the imaging devices used: a UI-3060CP-M-GL camera from IDS connected to a laptop computer, and a Raspberry Pi Camera V2 connected to a Raspberry Pi 3B+. We refer to the version with the IDS camera as the ‘IDS lab-prototype’ and the Raspberry Pi version as the ‘Raspberry Pi lab-prototype’.

The IDS camera has a sensor size of 11.345 mm × 7.126 mm with pixel size 5.86 × 5.86 *µ*m. The Raspberry Pi camera has a sensor of dimensions 3.68 × 2.76 mm with pixel size 1.12 × 1.12 *µ*m. The larger pixel size of the IDS camera provides a better signal to noise ratio and sensitivity. Due to sensor dimensions the IDS camera system had a rectangular field of view corresponding to a 200×100 *µ*m region within the sample, with the raspberry pi camera imaging a 130×95 *µ*m region in the sample.

The IDS camera was placed at the tube length of ≈160 mm from the focal plane producing a sub-Nyquist sampling rate (Nyquist sampling defined here as having pixel size of half the dimensions of the smallest resolvable features). Due to the small sensor size on the the Raspberry Pi camera, it was also operating sub-Nyquist sampling as it was placed much closer to the objective in order to obtain usable, visible images.

When using the lab-prototypes, images were captured at each plane by manually moving the sample using the Thorlabs translation stage with graduated micrometer with resolution of 10 *µ*m. For the water microscope a single stepper motor (a MikroElektronika 28BYJ-48) was attached on the z translation adjustment gear to allow accurate control of the z translation movement. This was controlled using the Raspberry Pi and an Arduino with an attached Motor Shield. The x and y control was carried out by hand using the two additional gears.

Illumination was controlled using python code running on attached Raspberry Pi 3B+ computers for both the lab-prototypes and the water scope. Image acquisition with the IDS camera used proprietary IDS software connected to a laptop computer running Windows 7. Image acquisition with the Raspberry Pi lab-prototype and the water scope was carried out using python running on the Raspberry Pi 3B+. Processing of all sets of images was carried out with a python script, available in the supplementary material, and ImageJ [23].

Data from all versions of the device were analysed in the same manner. As the distance between out of focus planes, Δz, was greater than the diameter of the algal cells the RI map needs appropriate interpretation. Due to the difference in focal-plane and optical axis resolution, the map allows us to distinguish sub-cellular changes of refractive index in the focal plane, but the individual values arise from the mean refractive index of the entire cell along the optical axis at that location. Once a RI map was obtained from the TIE code the cells were sampled manually using the select tool in ImageJ. Twenty measurements of the refractive index from regions throughout the entire cell were made for each cell, analysed and then averaged to obtain an estimated RI value for that individual cell. Sixteen cells, eight from each of two strains, were analysed on the lab-prototypes, while six cells, three from each strain, were analysed with the water scope.

## 5 Results

The IDS lab-prototype images in figure 2 (a-c) show a bright-field image, a TIE phase image and a RI index map of a single *Thalassiosira pseudonana* cell (CCAP 1085/12). Figure 2 (d-f) show equivalent images for the Raspberry Pi lab-prototype. Images used to create the figures and table in this work, along with python code and example raw images are available on the Strathclyde file sharing site.

**Figure 2:**
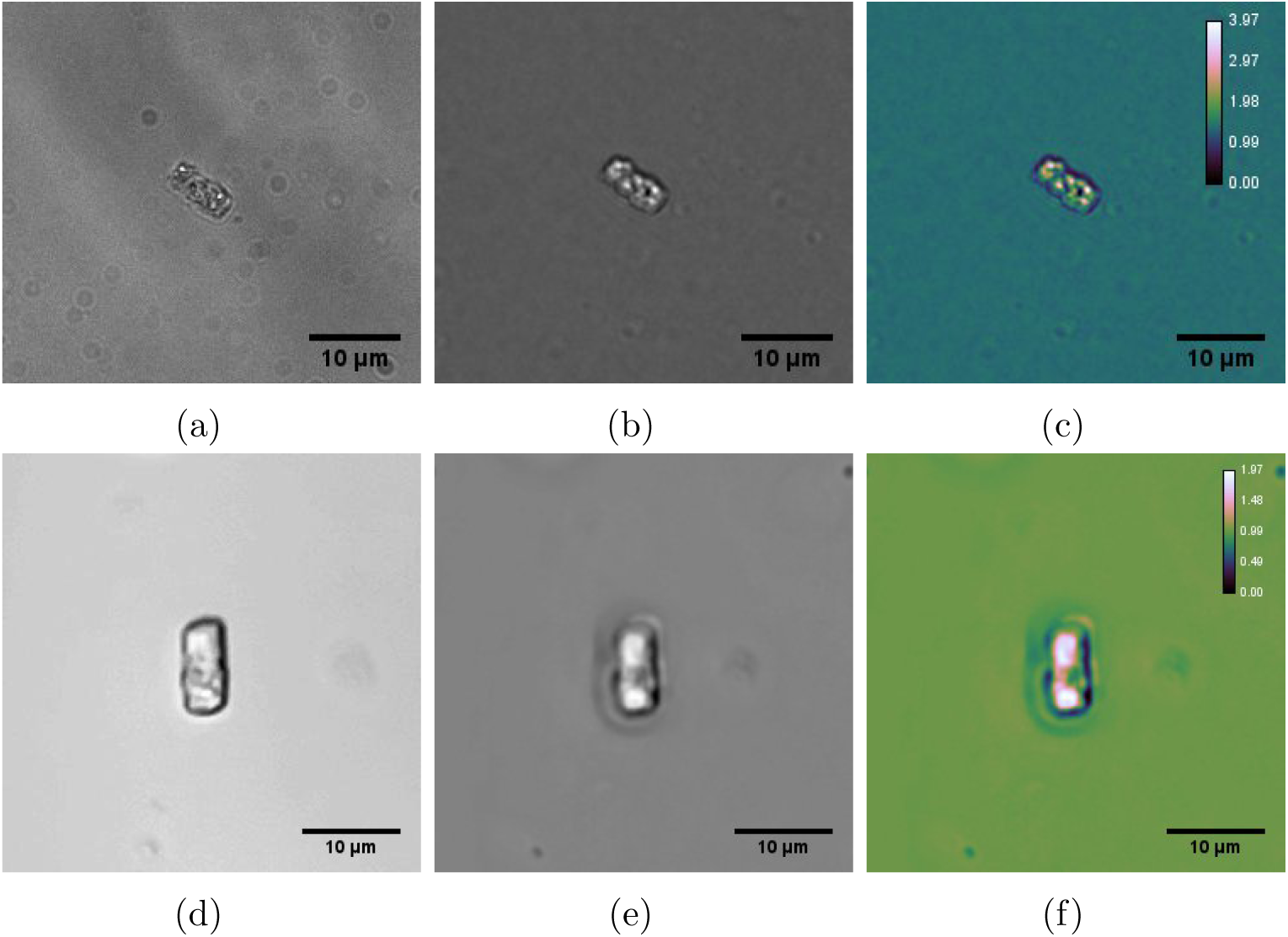
(a) Bright-field, focal plane image, (b) TIE image, and (c) RI map for *Thalassiosira pseudonana* captured with IDS camera. (d-e) Equivalent images captured with Raspberry Pi camera. The RI map is coloured using the Cube Helix Look Up Table [24]. This colourmap is designed to better illustrate increases in intensity by taking into account perceived brightness and is useful in illustrating the relative change in refractive index from the background water value.

Figure 3 shows the measured results for each of the instruments. Figure 3 (a) shows the twenty measurements taken with the IDS prototype for both 8x *T.pseudonana* and 8x *Emiliania huxleyi* (CCAP 920/12) cells. The violin plot on the right hand side shows the distribution of the measured refractive index values for the eight cells analysed. Figure 3 (b) and (c) show equivalent data for the Raspberry Pi lab-prototype and the Water scope respectively. The water scope data consists of 6 total cells as noted above.

**Figure 3:**
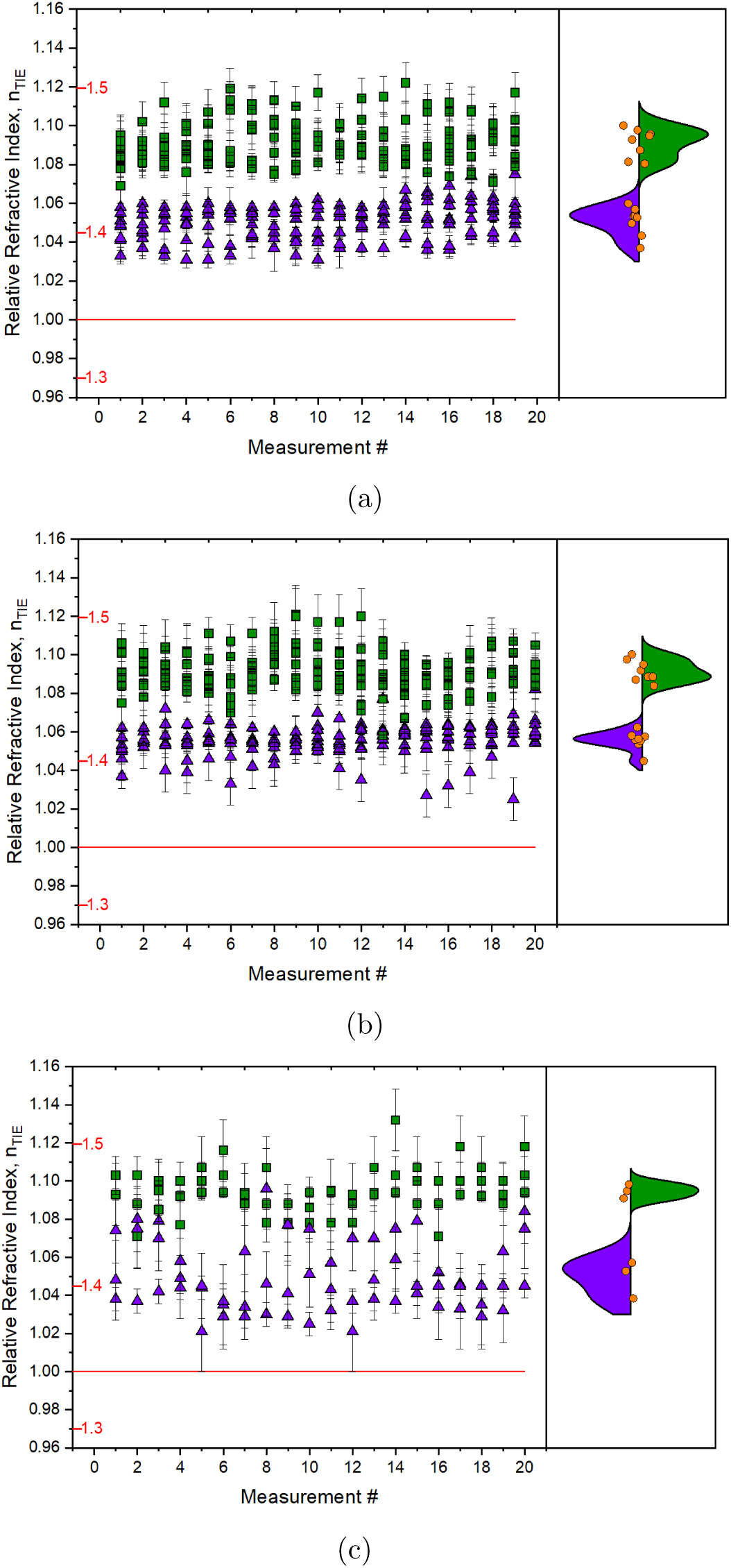
Comparison of measured samples taken with (a) IDS lab-prototype, (b) Raspberry Pi lab-prototype, and (c) water scope showing differences in measured refractive indices relative to water. *T.pseudonana* shown in purple triangles, *E.huxleyi* shown in green squares. The red tick marks show the absolute refractive index values. The orange circles in the right hand panel show the cell refractive index value for 8 *T.pseudonana* cells and 8 *E.huxleyi* cells (3 each for the water scope).

The refractive index values found are summarised in table 1. Using the IDS lab-prototype, the average refractive index for *T.pseudonana* was found to 1.051±0.007 relative to water, while the refractive index for *E.huxleyi* was found to be 1.091±0.006. Using the Raspberry Pi lab-prototype, the average refractive index for *T.pseudonana* was found to be 1.055±0.005 relative to water, while the refractive index for *E.huxleyi* was found to be 1.092±0.005. And finally, using the water scope, the average refractive index for *T.pseudonana* was found to be 1.049±0.008 relative to water and the refractive index for *E.huxleyi* was found to be 1.095±0.003.

**Table 1:** Mean refractive index for (a) *T.pseudonana* and (b) *E.huxleyi* obtained with IDS lab-prototype, (c) and (d) using the Raspberry Pi lab-prototype, and (e) and (f) using the water scope

| *Cell no.* | (a)<br>*T.pseudonana* | (b)<br>*E.huxleyi* | (c)<br>*T.pseudonana* | (d)<br>*E.huxleyi* | (e)<br>*T.pseudonana* | (f)<br>*E.huxleyi* |
| --- | --- | --- | --- | --- | --- | --- |
| 1 | 1.037 | 1.081 | 1.045 | 1.100 | 1.057 | 1.095 |
| 2 | 1.043 | 1.096 | 1.058 | 1.084 | 1.038 | 1.090 |
| 3 | 1.054 | 1.098 | 1.062 | 1.092 | 1.053 | 1.100 |
| 4 | 1.050 | 1.080 | 1.054 | 1.089 |  |  |
| 5 | 1.053 | 1.087 | 1.056 | 1.095 |  |  |
| 6 | 1.060 | 1.095 | 1.055 | 1.089 |  |  |
| 7 | 1.060 | 1.093 | 1.058 | 1.087 |  |  |
| 8 | 1.057 | 1.100 | 1.056 | 1.097 |  |  |
| average | 1.051 | 1.091 | 1.055 | 1.092 | 1.049 | 1.095 |
| std. dev | 0.007 | 0.006 | 0.005 | 0.005 | 0.008 | 0.003 |

## 6 Discussion

The refractive index values showed a clear distinction between the two species of algae tested. The values obtained (1.051 using the IDS camera and 1.055 using the Raspberry Pi camera) for *T.pseudonana* and (1.091 with the IDS camera and 1.092 with the Raspberry Pi camera) for *E.huxleyi*, were found to be in close agreement with published values obtained from alternative methods including flow cytometry, which reports on algal values from bulk water samples falling in the range 1.05-1.15 [25]. The refractive index may also be obtained by comparing the scattering efficiency Qb = Qc - Qa with theoretical scattering models. Reported values for this method @ 660 nm give values of 1.035-1.063 for *T.pseudonana* and 1.093-1.095 for *E.huxleyi* [26]. As noted in [27], coccolithophores typically have higher refractive index values than those of green algae due to their calcium carbonate shell. It is also noted that water content, metabolic condition, environmental variables, cell size, and cell age can all affect the optical properties, in particular the refractive index, of algal cells. These factors were not closely controlled in this work but information about some of them can be gleaned from information collected using this method. Cell size can be observed directly with the microscope while environmental variables could be monitored/controlled with the addition of temperature control hardware, illumination control etc. The addition of more robust and complete control of these factors presents an avenue for improvement and further work.

The Raspberry Pi camera, performing with pixel sizes greater than that required for Nyquist sampling, gave good agreement (within ±1 %) with the more expensive and sensitive IDS equipment. Comparison of the variances in the IDS data versus the Raspberry Pi data using a two-sample F-Test indicated no statistically significant difference between the methods implying comparable levels of precision. This suggests a low cost device as proposed in the introduction could be feasible in providing useful, actionable data to allow local communities in developing nations to discriminate between types of algae, if not on a species specific level then at a higher taxonomic level, and monitor increases in algal populations enabling precautionary action to minimise the risk of HABs developing, rather than reaction afterwards to mitigate the damage. The whole microscope and imaging hardware can fit in a volume of < 1000 cm^3^, making it portable, cheap and robust.

We have found that the TIE-based measurement approach reliably gives a narrow range of refractive index values for differing algae. However we also suggest that this is not sufficient by itself for unambiguous determination of an algal species from RI data alone. The quality of imaging obtained from these compact microscopes, combined with the potential to incorporate fluorescence-based imaged using high-power LED’s may provide a path to using multiple optical analytic techniques to narrow down potential candidates to give a warning ahead of HAB events.

The TIE allows for obtaining phase information without the need for expensive or complicated equipment. The ability to extract refractive index information provides an advantage over traditional phase microscopy methods while offering practical qualitative data from microscopy imagery. The limitations of the measurements include susceptibility to misalignment of the apparatus and assumptions regarding the background medium. These issues can potentially be overcome with the use of a 3D printed single part apparatus which fixes all optical elements in the correct position with little to no motion allowed reducing the potential for misalignment (while also reducing the cost of components), and the use of blank media or a medium of accurately known refractive index.

Taken as a relative measurement this method potentially allows for discrimination of objects based on their effect on refractive index against an unknown background value. Therefore, where differences in refractive index are sufficient, this low-cost, open-source, portable device may provide a useful tool, not only in relation to HABs but also in providing information regarding marine particles in general. Further development in terms of analytical capability holds the potential for enabling near real-time species identification and physiological assessment.

## Supplementary material

Supplementary material can be found on the associated Strathclyde file sharing site.

## Author contribution statement

S.G. and K.R. carried out the experiments. H.B. provided appropriate algal samples. S.G. analysed the data. S.G. wrote the manuscript with contributions from H.B., D.M., and B.R.P. B.R.P. conceived of the work and supervised the project.

## Research ethics statement

The work in this paper does not require this statement

## Animal ethics statement

The work in this paper does not require this statement

## Permission to carry out fieldwork statement

The work in this paper does not require this statement

